# Characterization of five purine riboswitches in cellular and cell-free expression systems

**DOI:** 10.1101/2021.04.14.439898

**Authors:** Milca Rachel da Costa Ribeiro Lins, Graciely Gomes Correa, Laura Araujo da Silva Amorim, Rafael Augusto Lopes Franco, Nathan Vinicius Ribeiro, Victor Nunes de Jesus, Danielle Biscaro Pedrolli

## Abstract

*Bacillus subtilis* employs five purine riboswitches for the control of purine *de novo* synthesis and transport at the transcription level. All of them are formed by a structurally conserved aptamer, and a variable expression platform harboring a rho-independent transcription terminator. In this study, we characterized all five purine riboswitches under the context of active gene expression processes both *in vitro* and *in vivo*. We identified transcription pause sites located in the expression platform upstream of the terminator of each riboswitch. Moreover, we defined a correlation between *in vitro* transcription readthrough and *in vivo* gene expression. Our *in vitro* assay demonstrated that the riboswitches operate in the micromolar range of concentration for the cognate metabolite. Our *in vivo* assay showed the dynamics of control of gene expression by each riboswitch. This study deepens the knowledge of the regulatory mechanism of purine riboswitches.

## INTRODUCTION

*B. subtilis* tightly controls its purine intracellular concentration employing different layers of regulation. Expression of the biosynthetic genes and genes coding for purine transporters (*pur* operon, *xpt*-*pbuX, pbuG, nupG*, and *pbuE*) is under control of purine riboswitches. Additionally, the *pur* operon, *xpt*-*pbuX*, and the *pbuG* gene are under control of the PurR repressor [1]. Permeases PbuX, PbuG, and NupG import purines from the environment when they are available. On the other hand, PbuE is an efflux pump, that keeps the cellular purine balance by pumping out adenine and hypoxanthine [2] (Fig. 1). Riboswitches are regulatory RNAs located in the messenger RNA and composed of an aptamer and an expression platform. The aptamer senses small molecule or ion ligands and the platform controls gene expression [3]. Purine riboswitches have emerged as models for studying RNA structure and folding, and small molecule recognition. Its ∼60 nt in length aptamer is one of the smallest among riboswitches. Moreover, it is structured as a multi-helix with helical packing, which is one of the simplest structures among riboswitch aptamers. Finally, unlike other RNAs, purine aptamers tend to fold correctly in different conditions and do not multimerize [4]. Such studies have deepened our knowledge of how riboswitch aptamers function to efficiently regulate gene expression [5–9]. Nonetheless, there are still some aspects of riboswitches that have not been elucidated yet. The lack of nucleotide conservation in the expression platform as well as how the riboswitches function in the cellular context are still poorly understood [4].

**Figure 1.**
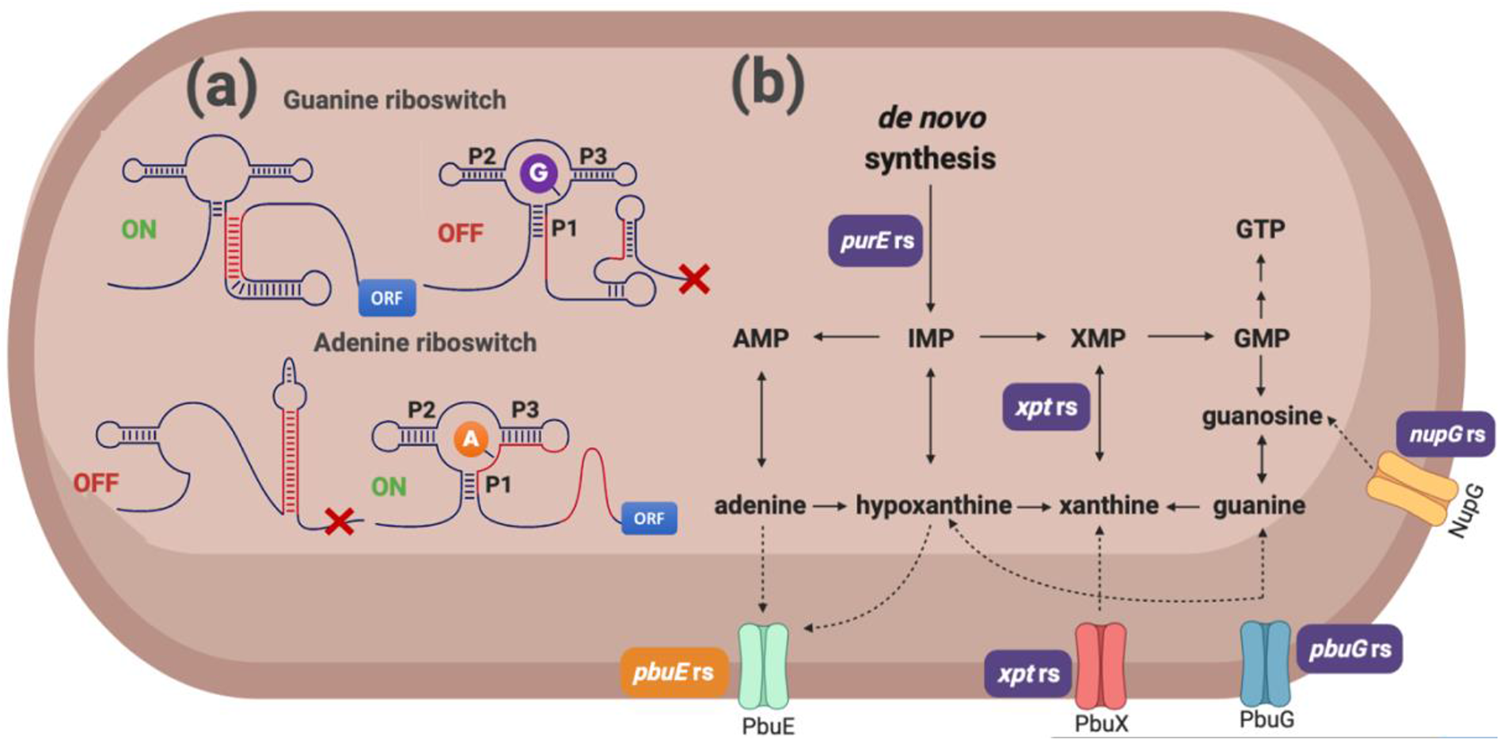
Control of the purine metabolism in *B. subtilis*. (A) Riboswitches *purE, xpt, nupG*, and *pbuG* turn OFF gene expression when bound to guanine. On the other hand, the *pbuE* riboswitch turns ON gene expression when bound to adenine. The riboswitches are named after the gene found directly downstream in the chromosome. The riboswitch aptamer folds into P1, P2, and P3 stems. (B) The purine riboswitches control the expression of genes active on purine transport and *de novo* synthesis. Purple boxes identified as *purE* rs, *xpt* rs, *nupG* rs, *pbuG* rs, and the orange box identified as *pbuE* rs indicate the metabolic process (transport and reaction catalysis) regulated by each riboswitch at the transcription level. Figure created in BioRender.com

Purine riboswitches specifically bind either guanine or adenine. In *B. subtilis*, guanine riboswitches *purE, xpt, nupG*, and *pbuG* turn OFF gene expression when bound to guanine. On the other hand, the adenine riboswitch *pbuE* turns ON gene expression when bound to adenine (Fig. 1). Conserved nucleotides are mostly found in the non-paired regions of the aptamer (Fig. 2a). Aptamers fold into a highly conserved structure constituted of P1 stem, and P2 and P3 stem-loops. Although structurally conserved, the stems have different lengths, resulting in different free energies (Fig. 2b). These features apply for both guanine and adenine aptamers. The only determinant for discrimination between the two purines is an unpaired cytosine or uracil in the stretch between P3 and P1, which forms a Watson-Crick pairing with the cognate purine [5, 7, 9–12]. On the other hand, the expression platform is not conserved at the nucleotide level. Although not conserved, all expression platforms are able to fold into a rho - independent terminator, responsible for the premature transcription termination determined by the aptamer status (bound or unbound) [9, 10, 12].

**Figure 2.**
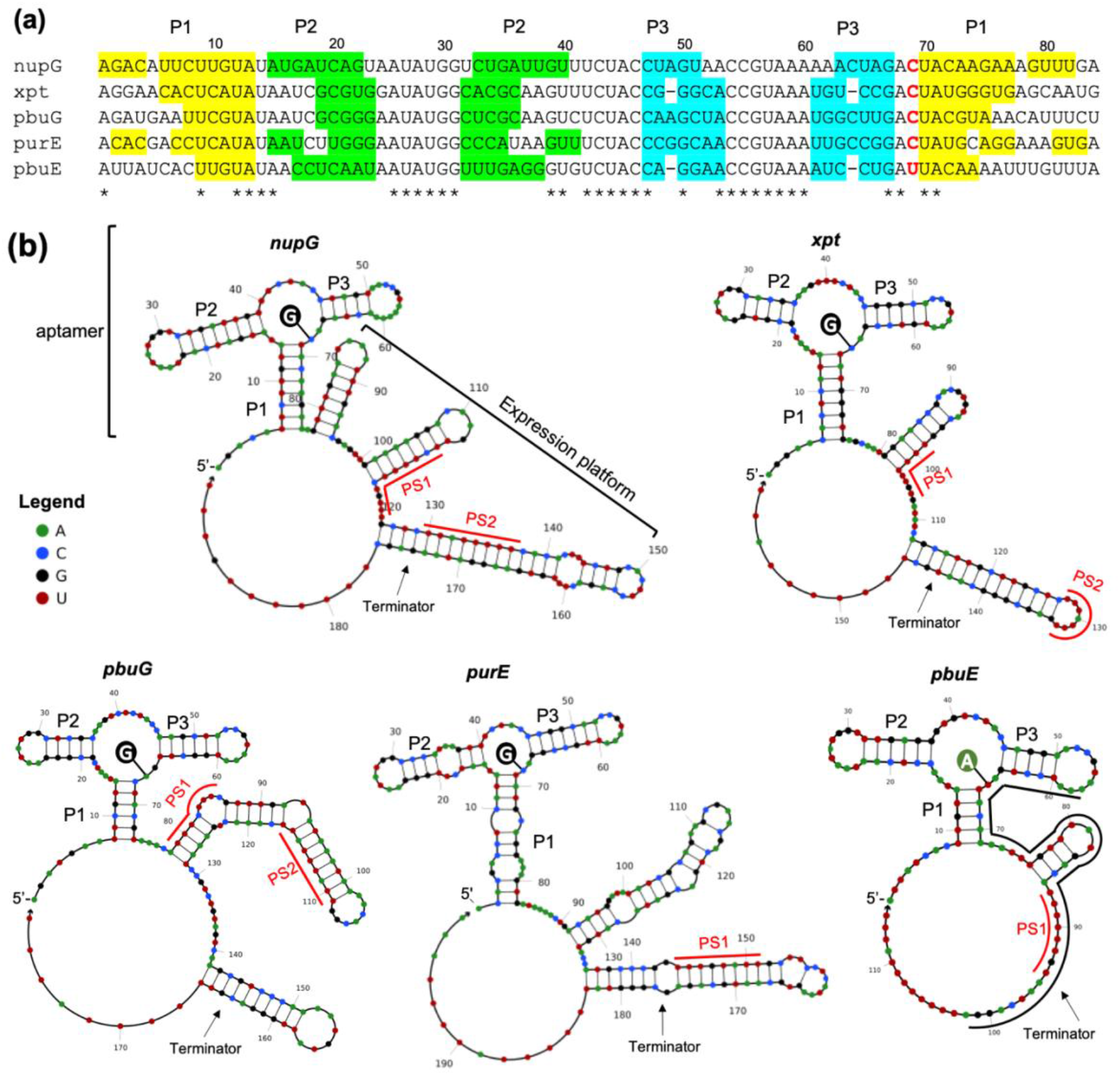
Sequence and folding of the five purine riboswitches found in *B. subtilis*. (a) alignment of the mRNA sequence domains corresponding to the riboswitch aptamers. Color-shaded ribonucleotides identify the characteristic aptamer stems P1, P2, and P3. The red-labeled base between P3 and P1 is responsible for discrimination between the ligands guanine (C) and adenine (U). (b) Secondary structure formed by the riboswitch aptamers and expression platforms when bound to the cognate purine. Structures represent the OFF state for the riboswitches *nupG, xpt, pbuG*, and *purE*, and the ON state for *pbuE*. Mapped transcription pause sites are labeled in expression platform as PS1 and PS2 (see figure 3). Structures generated with NUPACK Web Application.

The characterization of the purine riboswitches under the context of an active gene expression process, *in vitro* and *in vivo*, is greatly relevant for understanding their role in the cell. In this study, we characterized the five *B. subtilis* purine riboswitches under the context of active transcription and translation processes, both *in vitro* and *in vivo*, isolated from other gene expression regulatory processes.

## MATERIAL AND METHODS

### Sequences and cloning

Purine riboswitches *purE, xpt, nupG, pbuG*, and *pbuE* were amplified by PCR with Phusion High-Fidelity DNA Polymerase (Invitrogen) using genomic DNA of *B. subtilis* 168 as a template. Oligonucleotides were designed to contain restriction sites for *Bam*HI and *Hind*III, which were used to clone the PCR products into the P_*ribB*_-*luc* plasmid [13] using the same restriction sites. Each riboswitch was cloned upstream of the firefly luciferase gene (*luc*) and downstream of the *rib* promoter (P_*ribB*_) in the P_*ribB*_-*luc* plasmid. *E. coli* TOP10 was used for plasmid cloning and propagation. *E. coli* was cultivated in Lysogeny Broth (LB) enriched with 100 μg mL^-1^ ampicillin, and incubated in orbital shaker at 37°C and 220 rpm. Positive clones were confirmed by plasmid sequencing (3130 Genetic Analyzer, Applied Biosystems). Resulting sequences were analyzed using Sequence Scanner v 2.0 (Applied Biosystems). Another round of PCR was performed to amplify the *srfA* promoter and the purine riboswitches *purE, xpt, nupG, pbuG* and *pbuE* using genomic DNA of *B. subtilis* 168 as a template. Oligonucleotides for the P*srfA* were designed to contain restriction sites for *Eco*RI and *Xba*I, and oligonucleotides for the riboswitches were designed to carry the *Spe*I and *Pst*I restriction sites. The P_*srfA*_ and the riboswitch PCR products were digested and ligated to the pBS3C*lux* plasmid [14], previously digested with *Eco*RI and *Pst*I, to generate plasmids pBS3C-P_*srfA*_-riboswitch-*lux*. A control plasmid carrying the P_*srfA*_ but no riboswitch was also constructed by ligating *Eco*RI and *Xba*I digested P_*srfA*_ PCR to *Eco*RI and *Spe*I digested plasmid. After cloning confirmation by sequencing, each plasmid, carrying one purine riboswitch, was separately used to transform in *B. subtilis* 168. Integration of each plasmid into the genomic *sacA* locus was confirmed by PCR. All DNA sequences, plasmids, and strains used are listed in the Supplementary material (Tables S1, S2 and S3).

### *In vitro* transcription

Linear DNA templates for *in vitro* transcription were prepared by PCR using Phusion High-Fidelity DNA Polymerase (Invitrogen) and the P_*ribB*_-riboswitch-*luc* plasmid series. The T7 promoter and a 36 nt spacer were added to the forward primers, and the reverse primers were designed to amplify 164 nt of the *luc* gene. Five linear template sequences were PCR amplified corresponding to the five purine riboswitches. PCR products were than extracted from agarose gels and purified using the Wizard^®^ SV Gel and PCR Clean-Up System (Promega). Eluted DNA solutions were then precipitated with ethanol and resuspended in RNase-free water.

RNAs transcripts were generated by *in vitro* transcription using the RiboMAX^™^ Large Scale RNA Production Systems (Promega). Reactions were set up by adding 2 µL of the T7 Transcription 5X Buffer, 2 µL rNTP mix to either 5 mM or 150 µM final concentration, 200 ng of PCR templates, 1 µL T7 enzyme mix, and RNase-free water up to 10 µL. Reaction mixes were incubated at 37°C for up to 1h in a dry bath. Inactivation was performed at 70°C for 20 min. To eliminate the template, all samples were treated with RQ1 RNase-Free DNase (Promega) for 3 hours at 37°C. Samples were then precipitated with ethanol and resuspended in the same volume with RNase-free water.

### High-resolution electrophoresis

*In vitro* transcribed RNAs were analyzed by high resolution electrophoresis using the RNA 6000 Nano Kit (Agilent). Purified RNA transcript solutions were incubated at 95 °C for 2 min for denaturation. 1 µL of heat denatured sample was loaded on the chip. Electrophoresis was developed in the 2100 Bioanalyzer system (Agilent) according to the manufacturer instructions.

### *In vitro* gene expression

The regulatory activity of the five purine riboswitches was measured by *in vitro* gene expression reaction in the presence and absence of guanine or adenine. Each P_*ribB*_-riboswitch-*luc* plasmid was extracted from overnight grown *E. coli*, and purified twice with a miniprep kit (PureYield^™^ Plasmid Miniprep System, Promega). Plasmids were eluted with RNase free water and used as templates for *in vitro* gene expression (*E. coli* S30 Extract System for Circular DNA, Promega).

Reactions were set up by adding 1 μL of 0.25 mM amino acid mix, 4 μL of S30 Premix, 3 μL of S30 Extract, 1 μL of purine stock solution, and 1 μL of plasmid (final concentration 10 ng μL^-1^). Guanine (Merck G11950) and adenine (Merck A8626) were added to the reactions at 50, 100 or 250 μM. Ten times concentrated purine stock solutions were prepared in 20 mM Tris-HCl buffer pH 8.0. To ensure solubility, each purine was first solubilized in 100 mM HCl solution, which was then used to prepare the 20 mM Tris-HCl purine solution. The same buffer was used in the control reactions without purine. The reaction mixture was incubated at 30°C for 20 min, and 90 μL of 1% (m/v) bovine serum albumin was added to stop the reactions. Each treatment was performed in triplicate.

To quantify the luciferase activity, 10 μL of stopped reactions were added to 50 µL of luciferase substrate solution (Luciferase Assay System kit, Promega) in a 96-well white microplate. Luminescence readings were taken on a microplate reader (Tecan Infinite 200 Pro, Switzerland). All luciferase activities were normalized to the activity level measured for no purine treatment and named as fold change (FC).

### *B. subtilis* growth and gene expression

*B. subtilis* was cultivated in Luria-Bertani (LB) broth or mineral medium (MM: 4 g L^−1^ KH_2_PO_4_, 16 g L^−1^ K_2_HPO_4_ 3H_2_O, 3.3 g L^−1^ (NH_4_)_2_SO_4_, 0.232 g L^−1^ MnSO_4_ 4H_2_O, 12.3 g L^−1^ MgSO_4_ 7H_2_O, 0,136 g L^−1^ ZnCl2, 0.05 g L^−1^ tryptophan, 0.022 g L^−1^ iron ferric ammonium citrate, 5 g L^−1^ glucose). 5 μg mL^-1^ chloramphenicol was added to the culture medium for the recombinant strains. Inocula were prepared from overnight cultures grown in an orbital shaker at 37°C and 220 rpm. Main cultures were inoculated to OD_600_ of 0.1 in 200 µL LB supplemented with 100 μM guanine or adenine, on white 96-well microplates with optically clear bottom. To prepare adenine and guanine enriched LB we first solubilized each purine in 1 mM HCl. Then we added the LB components and neutralized the pH with NaOH. All treatments were performed in biological triplicates. Luminescence and OD_600_ measurements were taken during growth.

### Luminescence and absorbance measurements

Luciferase activity measurements from *in vitro* gene expression samples were performed in white 96-well microplates with flat bottom incubated at 26°C. Luminescence emission was measured every 30 s for 10 min on a microplate reader (Tecan Infinite 200 Pro, Switzerland) set with 1,000 ms for integration time. Luminescence measurements from growing *B. subtilis* were carried out in white 96-well microplates with optically clear bottoms. Culture density was monitored by absorbance measurements at 600 nm (OD_600_) taken from the same transparent bottom microplates. Plates were incubated at 37°C and orbital shaking of 143 rpm. Measurements were performed every 10 min for five hours in the microplate reader.

### Riboswitch structure analysis

Riboswitch folding structures and free energy were generated using the analysis tool within the NUPACK Web Application [15].

## RESULTS

### Pause sites are check points for the aptamer status

All five purine riboswitches in *B. subtilis* control gene expression at the transcription level by alternative folding that generates a premature transcription termination [9, 10]. Such mechanism limits the regulatory activity to a short timeframe from the transcription start to the point right before the RNA polymerase escapes beyond the riboswitch transcription terminator. In that short timescale of transcription, riboswitches must rapidly and efficiently fold into the aptamer structure, survey the cellular environment for the cognate metabolite, and switch to the alternative secondary structure (or not) according to the aptamer status (bound or unbound) [4, 9, 16]. Transcription pause sites downstream of the aptamer and upstream of the transcription terminator have been described for the FMN and the *pbuE* riboswitches [9, 16, 17]. They are uracil rich sequences that withhold the RNA polymerase and provides extra time for ligand binding before the formation of the thermodynamically favored termination hairpin.

We used *in vitro* transcription to map the pause sites in the purine riboswitches. We first run the reactions using 5 mM rNTPs to mark the full-length and terminated RNA bands generated. Then we dropped the rNTP concentration to 150 µM. By limiting the rNTP supply we expected to increase the residence time of the RNA polymerase in the pause sites in order to map them. At high rNTP supply the *nupG* riboswitch is transcribed into a 220 nt-long terminated RNA, and into a 430 nt-long readthrough RNA. However, at limited rNTP supply transcription under control of the *nupG* riboswitch accumulates other two smaller bands corresponding to the pause site 1 (160 nt-long) and pause site 2 (175 nt-long) (Fig. 3a). These gel bands correspond to the U-rich sites labeled as PS1 and PS2 in the aptamer structure shown in Fig. 2b. Note that, for the *in vitro* transcription reaction we used a template carrying a 36 bp sequence downstream of the promoter and upstream of the riboswitch aptamer; therefore, the resulting RNA bands are longer than the numbering shown in Fig. 2. Similarly, the *xpt* and the *pbuG* riboswitches had two transcription pauses mapped (Fig. 3b and 3c; Fig. 2b). On the other hand, we mapped only one pause site for the *purE* and *pbuE* riboswitches (Fig. 3d and 3e; Fig. 2b). From all five riboswitches, only the *pbuE* had its pause site previously mapped [16]. It was latter predicted for the *pbuE* riboswitch two pause sites at U-rich sequences 74-80 and 87-95 [17]. We confirmed that the *pbuE* riboswitch has only one pause site at the 87-95 U-rich sequence (Fig. 2b).

**Figure 3.**
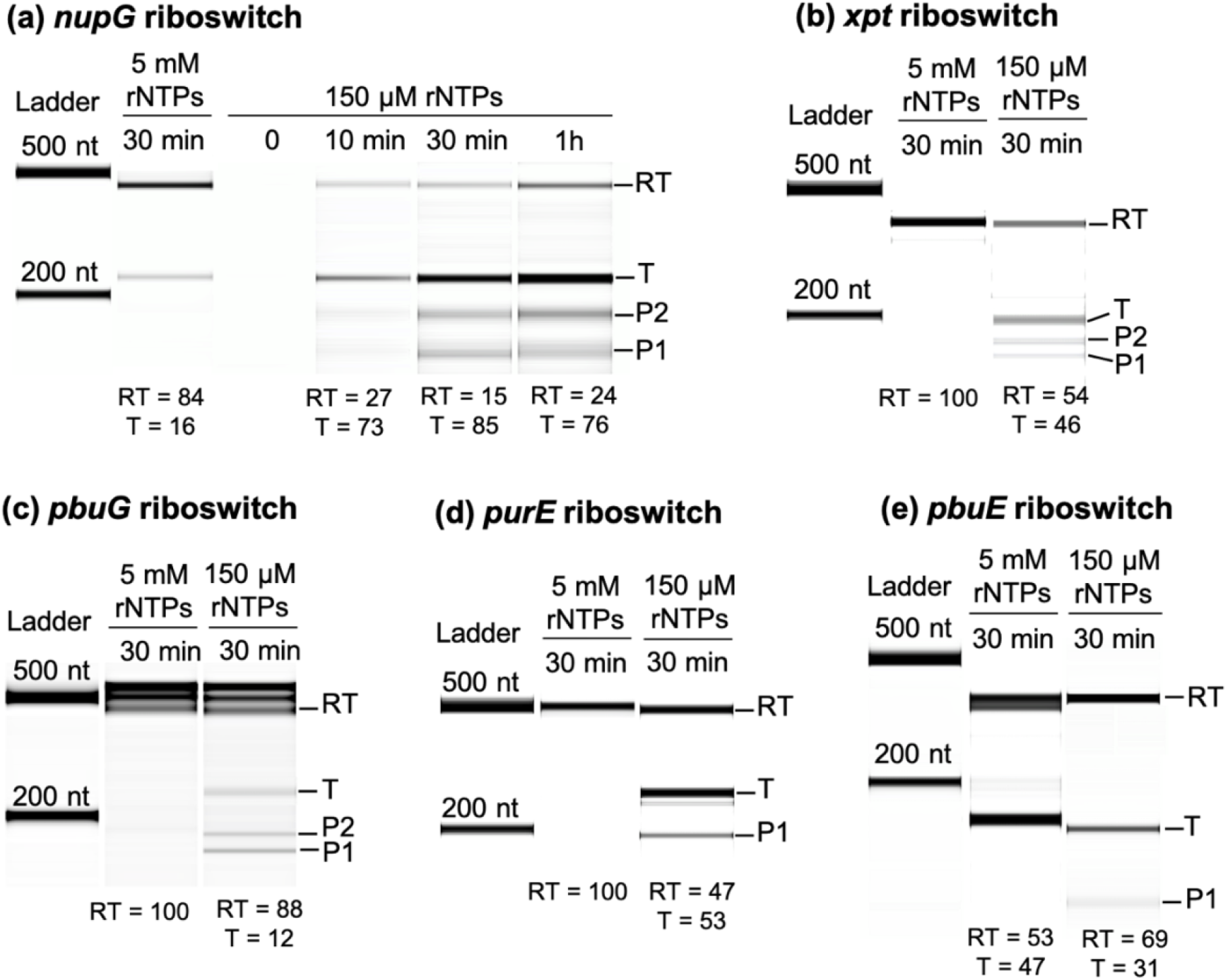
Mapping the transcription pause sites in the riboswitches’ expression platform. Riboswitches were transcribed *in vitro* using 5 mM or 150 µM rNTP in the reaction, which was incubated up to 1h. The resulting RNA products were resolved using high resolution electrophoresis and identified as readthrough (RT), terminated (T), pause site 1 (PS1), and pause site 2 (PS2). (a) *nupG* riboswitch. (b) *xpt* riboswitch. (c) *pbuG* riboswitch. (d) *purE* riboswitch. (e) *pbuE* riboswitch. In (c) the bands that appear to be larger than the readthrough RNA (RT) are folded RT molecules that resisted to the sample treatment at denaturing conditions. RT and T numbers below each lane refers to the percentage of readthrough and terminated RNAs.

### Control of gene expression *in vitro*

The five purine riboswitches *purE, xpt, nupG, pbuE*, and *pbuG*, were cloned between the P_*ribB*_ and the luciferase gene in the p P_*ribB*_-*luc* to generate pP_*ribB*_-riboswitch-*luc* plasmids. P_*ribB*_ is a constitutive *E. coli* promoter two orders of magnitude weaker than the T7 promoter (measured by *in vitro* gene expression, data not shown). Each plasmid was used as a template for *in vitro* gene expression in the presence of different concentrations of guanine and adenine. Reaction samples were assayed for luciferase activity, and the output was taken as a measurement for the riboswitch activity.

First, to validate our *in vitro* assay for riboswitches, we tested the *ribG* FMN riboswitch of *B. subtilis* cloned in the same pP_*ribB*_-*luc. ribG* promotes transcription termination upon FMN binding and has been characterized by *in vitro* gene expression elsewhere, but driven by the T7 promoter [18]. Treatment with 100 µM FMN caused a 4.6-fold decrease in gene expression under control of the *ribG* riboswitch, confirming the proper function of our assay. Next, we tested the purine riboswitches under the same conditions, but using adenine or guanine as additive. Results show that all five purine riboswitches discriminate between guanine and adenine, and regulate gene expression *in vitro* in a dose-dependent way (Fig. 4a-e). *purE, xpt, nupG* and *pbuG* riboswitches efficiently downregulate gene expression when treated with guanine in the micromolar range of concentration. At the highest concentration tested (250 µM) gene expression was repressed 4.4-fold by *purE*, 4.1-fold by *xpt*, 3.2-fold by *nupG*, and 3.3-fold by the *pbuG* riboswitch. Adenine had none or a very small effect on the three guanine riboswitches (Fig. 4a-d). Although structurally similar to the other purine riboswitches, the *pbuE* riboswitch employs a different mechanism, it upregulates gene expression upon adenine binding. Like the guanine riboswitches *pbuE* respond to the cognate metabolite in the micromolar range, activating gene expression 5.7-fold at 250 µM adenine (Fig. 4e). Guanine had a much smaller activating effect on the *pbuE* riboswitch. *luc* expression from the control template pP_*ribB*_-*luc* is not affected by guanine or adenine (Fig. 4f).

**Figure 4.**
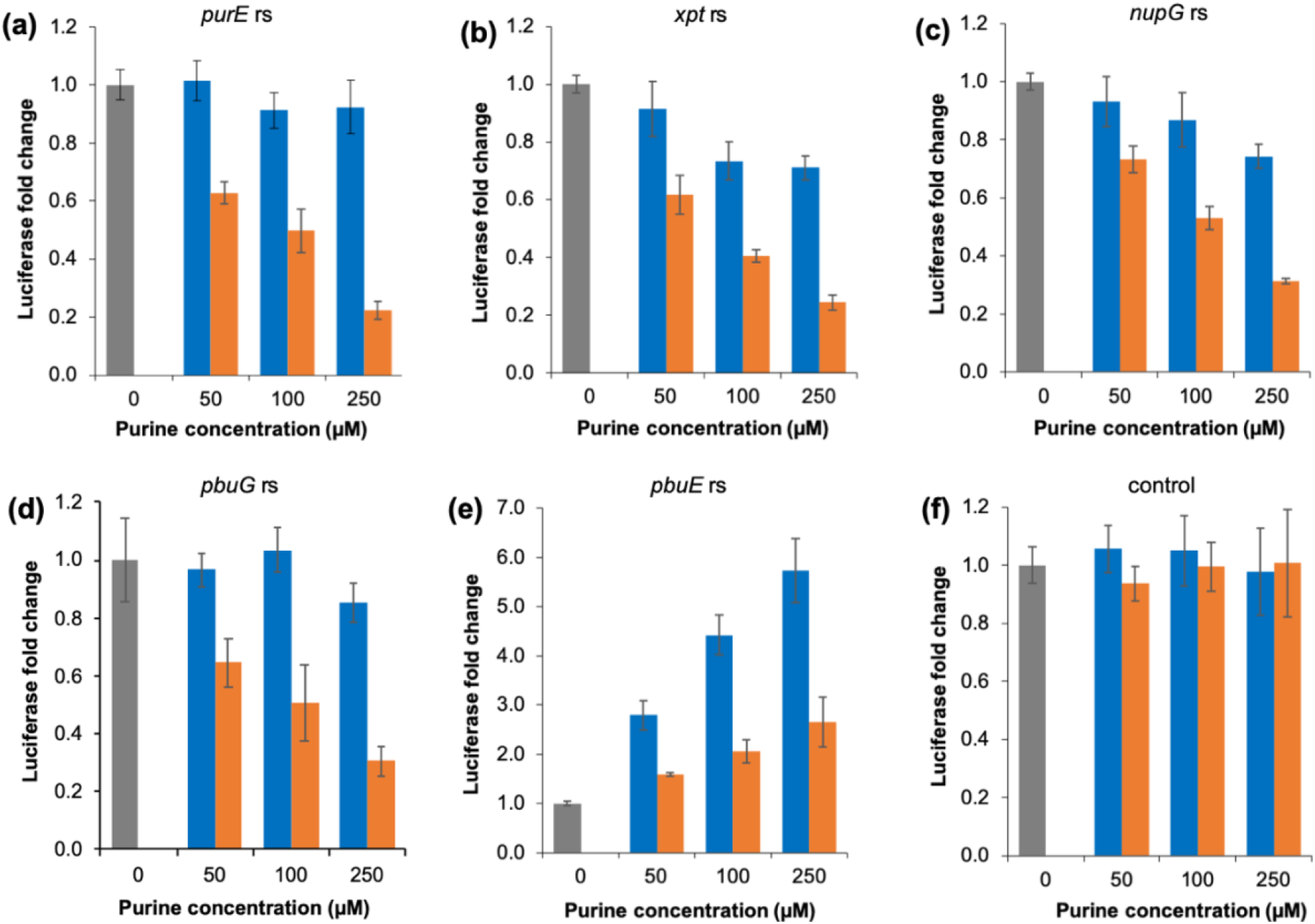
Regulatory activity of the purine riboswitches measured *in vitro*. Guanine (orange bars) or adenine (blue bars) were supplied to the *in vitro* gene expression reactions in different concentrations. All templates tested carry the firefly luciferase gene under control of the *ribB* promoter and the (a) *purE* riboswitch; (b) *xpt* riboswitch; (c) *nupG* riboswitch; (d) *pbuG* riboswitch; or the (e) *pbuE* riboswitch. (f) control without riboswitch. Firefly luciferase activity, measured from the *in vitro* gene expression reactions, was taken as a measure for the regulatory activity of the riboswitch. Luciferase activities were normalized to the activity level measured for no purine treatment. Note that the luciferase fold change scale is different for (d). Error bars represent the standard deviation calculated from three independent reactions.

### Apparent ligand affinity

The amount of purine needed for a 50% reduction (T_50_) of the luciferase activity in the *in vitro* assay was taken as a measure for the apparent ligand affinity of the riboswitch aptamers in the context of an active gene expression process. For guanine riboswitches, T_50_ was estimated by fitting the plot of the guanine concentration versus the inverse of the fold change (1/FC). For the adenine riboswitch, T_50_ was estimated by fitting the plot of the adenine concentration versus the fold change (Fig. S1). The lowest T50 was estimated for *pbuE* as 52 µM, followed by *xpt* with 76 µM, *purE* with 80 µM, and *pbuG* with 105 µM. The highest T_50_ was estimated for *nupG* as 115 µM.

*In vitro* gene expression assays were carried out in order to characterize the isolated function of the purine riboswitches. The next set of experiments *in vivo* revealed the interaction of the regulatory activities of the riboswitches with the purine uptake and metabolism in the cell.

### Control of gene expression *in vivo*

All five *B. subtilis* purine riboswitches were cloned into the pBS3C*lux* plasmid [14], together with the P_*srfA*_ promoter, in order to control the expression of the *luxABCDE* operon. Each pBS3C-P_*srfA*_-riboswitch-*lux* plasmid was integrated into the *B. subtilis* genome at the *sacA* locus. The resulting strains were cultivated in the presence of guanine or adenine, or without purine additive. Two riboswitches, *purE* and *pbuE*, were first tested in mineral medium (MM) and in LB to determine the influence of the purine content of LB in the riboswitch activity. No significant interference was found, and both riboswitches performed similarly in both media when treated with 100 µM of purine (Fig. S2). LB was chosen for further test because unlike MM no growth improvement was detected for the treatments compared to the control (LB only). It is important to say that the host *B. subtilis* 168 is prototrophic for both adenine and guanine; thus, purine supplementation is not required for growth. Measurements of OD_600_ and luminescence were taken during the course of culture growth. All engineered strains grew as well as the *B. subtilis* wt, no matter the treatment applied (Fig. S3). All riboswitches displayed regulatory control over the *lux* operon expression when exposed to the cognate metabolite, guanine or adenine (Fig. 5a-e). The control strain, whose *lux* operon is not controlled by a riboswitch, showed a similar luminescence production pattern for all three treatments applied (Fig. 5f). For all five riboswitches a regulatory effect was observed after 2h of cultivation, when the cultures entered the log growth phase (Fig. 5 and Fig. S3). Guanine riboswitches *purE, xpt, nupG*, and *pbuG* decreased 1.7 to 2.4-fold gene expression after 4h of treatment with guanine, compared to no purine treatment (Fig. 5a-d). The tightest control among the guanine riboswitches was exerted by the *pbuG* riboswitch from 3.2 to 3.6h of growth, when gene expression decreased 5.2-fold. Interestingly, adenine had no effect on the *nupG* riboswitch (Fig. 5c). When treated with adenine *purE* and *pbuG* riboswitches decreased gene expression during the log growth phase. However, *purE, pbuG*, and *xpt* riboswitches increased gene expression during the transition from log to stationary phase (4-5h) when treated with adenine. This increasing effect has not been observed in the *in vitro* gene expression assay. The adenine riboswitch *pbuE* showed a marked increase in gene expression after 3.5h of treatment with adenine, reaching 2.4-fold increase after 4.8h compared to the control. When compared to the guanine treatment, adenine supplementation caused a 7-fold increase in gene expression after 4.8h (Fig. 5e). Interestingly, a 4-fold suppression of gene expression was measured when guanine was added to the culture medium compared to the control. This strong guanine suppression effect has not been anticipated by the *in vitro* gene expression.

**Figure 5.**
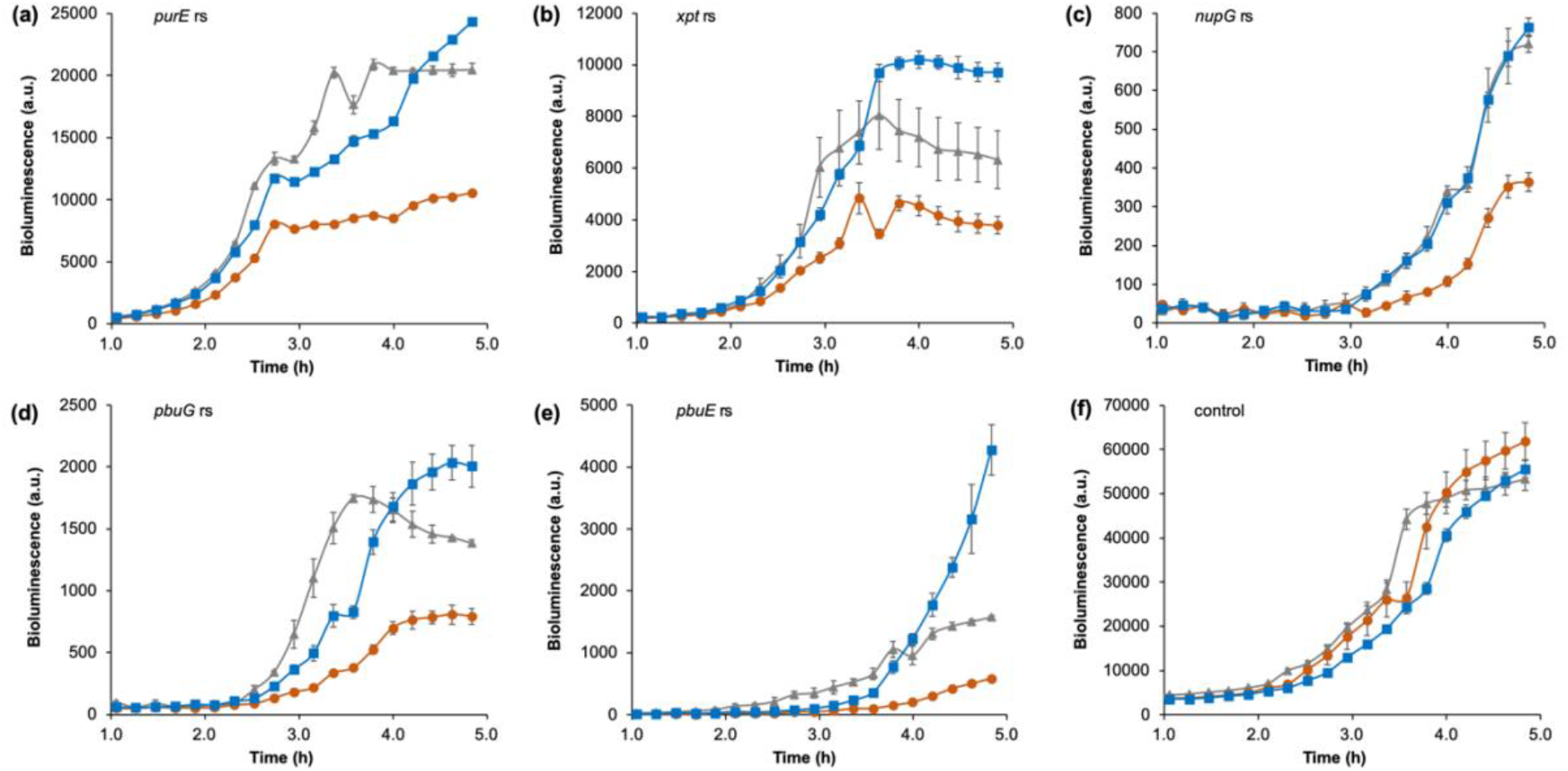
Regulatory activity of the purine riboswitches measured in the time-course of *B. subtilis* growth. Three treatments were performed: control - LB only (gray triangles), LB + 100 µM guanine (orange circles), and LB + 100 µM adenine (blue squares). Bioluminescence measured from *B. subtilis* cultures was taken as a measure for the regulatory activity of the riboswitches. All strains tested carry one genomic copy of *luxABCDE* under control of the (a) *purE* riboswitch; (b) *xpt* riboswitch; (c) *nupG* riboswitch; (d) *pbuG* riboswitch; or the (e) *pbuE* riboswitch. (f) control strain carrying one genomic copy of *luxABCDE* not regulated by a riboswitch. Note that the bioluminescence scales are different for each graph. Error bars represent the standard deviation calculated from biological triplicates.

Noteworthy, the riboswitches had a strong influence on the levels of gene expression reached. Compared to the levels achieved for the control (without riboswitch), the *purE* riboswitch reduced gene expression 2.5-fold, *xpt* caused 6.6-fold decrease, *pbuE* caused 12.5-fold decrease (adenine-induced level), *pbuG* 30-fold decrease, and the *nupG* riboswitch weakened the most gene expression decreasing it 74-fold (Fig. 5 and Fig. S4). Our reporter unit carries the same promoter, RBS, and reporter operon for all tested riboswitches, meaning the decreasing effect on gene expression could only be caused by the riboswitch. Indeed, there is a clear correlation between the transcription readthrough and the maximal measured reporter expression for all five riboswitches (Fig. S5).

## DISCUSSION

Since discovery riboswitches have been studied extensively using *in vitro* approaches, mostly employing RNA aptamer-metabolite binding assays to access affinities and dissociation constants (*K*_*D*_). Guanine riboswitch aptamers had their *K*_*D*_ for the cognate metabolite estimated in the low nanomolar range [10, 19], and the adenine riboswitch aptamer had its *K*_*D*_ estimated around 300 nM for 2-aminopurine (a fluorescent adenine analog) [5, 8, 9]. However, equilibrium between the aptamer and its cognate metabolite can only be reached in a timeframe longer than the time needed for the RNA polymerase to escape beyond the riboswitch transcription terminator. Transcription pause sites in the expression platform extend the timeframe for the metabolite-aptamer binding. Interestingly, carrying one or two pause sites has no effect on the apparent aptamer-ligand affinity or in the level of gene expression controlled by the riboswitches. Recently, transcription and translation in *B. subtilis* were described as uncoupled processes [20], which possibly confers some extra time for the metabolite-aptamer binding. Still, the timeframe may not be enough for equilibrium to be reached. The lifetime of the transcription pauses has been estimated around 60 s for both the *ribD* (FMN) and the *pubE* (adenine) riboswitches [16, 21]. Therefore, it has been proposed that transcriptionally regulated riboswitches are kinetically driven rather than thermodynamically controlled [9, 16]. To fit the kinetic model of control, the metabolite concentration required to elicit half maximal regulatory response (EC_50_ for *in vivo* and T_50_ for *in vitro*) must be greater than the *K*_*D*_ [4]. Our *in vitro* gene expression assay indicated T_50_ from 52 to 115 µM for the *B. subtilis* purine riboswitches, which is about 160-fold greater than the *K*_*D*_ for *pbuE*, and at least 10,000-fold greater than the *K*_*D*_ estimated for the *B. subtilis* guanine riboswitches. The estimated T_50_ for the purine riboswitches are in line with the T_50_ of 46 µM found for the *ribG* FMN riboswitch of *B. subtilis* [18]. The *ribG* riboswitch also had its *K*_*D*_ estimated in the low nanomolar range [9].

We tested all five riboswitches as genomic integrated single copy, and under the same genetic context (i.e. driven by the same promoter, and controlling the same reporter gene); therefore, we were able to measure the regulatory activity displayed by each of them only, without interference of promoter strength or the PurR repressor. All purine riboswitches control gene expression with dynamic ranges between 1.7 to 5.2. In this regard, our *in vivo* assay closely reproduced the fold changes achieved *in vitro*, demonstrating that no cellular component is required other than the transcription and translation machinery (and salts) to support the folding and the regulatory activity of purine riboswitches. In exponentially growing *E. coli* (glucose-fed) guanine and adenine are found in micromolar concentrations (guanine = 190 µM; adenine =1.5 µM) [22]. If that is also true for *B. subtilis*, any regulatory process that directly responds to these purines might operate in the same concentration range. There is though one study indicating that *B. subtilis* intracellular level of guanine ranges from 20 to 35 µM [2]; however, the nutritional condition and the culture growth phase for the test are not described. Our *in vitro* and *in vivo* assays on the regulatory activity demonstrated that the purine riboswitches of *B. subtilis* indeed respond to micromolar levels of guanine or adenine. Sensitivity in the micromolar range has been previously demonstrated for the *xpt* [23] and the *pbuE* [8] riboswitches in a cellular context. However, in both studies an amplification strategy was used, either as a regulatory cascade [23] or employing self-replicating plasmid [8]. Another study accessed the regulatory activities of the *purE, xpt* and *nupG* (former *yxjA*) riboswitches through RT-qPCR of endogenous transcripts [10]. Surprisingly, guanine-induced transcription termination was only detected when cells were treated with a metabolite concentration as high as 1.8 mM.

Previous studies on the regulatory function of riboswitches *in vivo* show end-point reporter results only [2, 8, 10, 23]; therefore, missing information about the temporal dynamics of the riboswitch regulation. Our results show that the regulatory activity of riboswitches depends on the culture development. For most treatments, gene expression regulated by riboswitches reached a plateau during the transition to the stationary phase, reaching steady state. That is possibly due to the reduced promoter activity during this growth phase. Noteworthy, the *pbuE* riboswitch starts to react late and keeps increasing gene expression during the transition to the stationary phase. Treating guanine riboswitches with adenine resulted in expression levels similar to the control until the early log growth phase. However, in the late log and transition to the stationary phase, gene expression increased above the control. Conversely, treating the adenine riboswitch with guanine turned off gene expression bellow control levels. High intracellular guanine concentration, caused by supplementation, triggers the intrinsic *purE* riboswitch and causes premature transcription termination of the *pur* operon. Accordingly, the *de novo* synthesis of purines is turned off ceasing the major source of cellular adenine. On the other hand, the control relies on the *de novo* synthesis to increase the adenine pool, which in turn activates slightly the *pbuE* riboswitch.

This study has investigated the metabolite-dependent regulation of the five *B. subtilis* purine riboswitches in the context of active gene expression processes *in vitro* and *in vivo*. We mapped the transcription pause sites in the expression platforms, and we demonstrated how each riboswitch works isolated from other regulatory processes naturally associated with them in cells.

## SUPPLEMENTARY MATERIAL

**Figure S1**. Purine riboswitches apparent affinity (T_50_) determined by *in vitro* gene expression.

**Figure S2**. Purine riboswitches performance in *B. subtilis* cultivated in mineral medium (MM) or LB medium supplemented with guanine or adenine.

**Figure S3**. Growth of *B. subtilis* strains. OD_600_ measurements were taken during growth of *B. subtilis* strains.

**Figure S4**. Relative gene expression under control of the purine riboswitches in *B. subtilis*.

**Figure S5**. Correlation between the transcription readthrough and the measured reporter expression under control of the purine riboswitches.

**Table S1**. Plasmids used in this study

**Table S2**. DNA parts and oligonucleotides used in this study **Table S3**. Strains used and generated in this research **Sequences**

## REFERENCES

1. Johansen LE, Nygaard P, Lassen C, et al (2003) Definition of a Second Bacillus Subtilis Pur Regulon Comprising the Pur and xpt-pbuX Operons Plus pbuG, nupG (yxjA), and pbuE (ydhL). J Bacteriol 185:5200–5209. https://doi.org/10.1128/JB.185.17.5200-5209.2003

2. Nygaard P, Saxild HH (2005) The Purine Efflux Pump PbuE in Bacillus subtilis Modulates Expression of the PurR and G-Box (XptR) Regulons by Adjusting the Purine Base Pool Size. J Bacteriol 187:791–794. https://doi.org/10.1128/jb.187.2.791-794.2005

3. Winkler W, Nahvi A, Breaker RR (2002) Thiamine derivatives bind messenger RNAs directly to regulate bacterial gene expression. Nature 419:952–956. https://doi.org/10.1038/nature01145

4. Porter EB, Marcano-Velázquez JG, Batey RT (2014) The purine riboswitch as a model system for exploring RNA biology and chemistry. Biochim Biophys Acta 1839:919–930

5. Lemay J-F, Penedo JC, Tremblay R, et al (2006) Folding of the adenine riboswitch. Chem Biol 13:857–868. https://doi.org/10.1016/j.chembiol.2006.06.010

6. Mandal M, Breaker RR (2003) Adenine riboswitches and gene activation by disruption of a transcription terminator. Nat Struct Mol Biol 11:29–35. https://doi.org/10.1038/nsmb710

7. Batey RT, Gilbert SD, Montange RK (2004) Structure of a natural guanine-responsive riboswitch complexed with the metabolite hypoxanthine. Nature 432:411–415. https://doi.org/10.1038/nature03037

8. Marcano-Velázquez JG, Batey RT (2015) Structure-guided mutational analysis of gene regulation by the Bacillus subtilis pbuE adenine-responsive riboswitch in a cellular context. J Biol Chem 290:4464–4475. https://doi.org/10.1074/jbc.M114.613497

9. Wickiser JK, Cheah MT, Breaker RR, Crothers DM (2005) The kinetics of ligand binding by an adenine-sensing riboswitch. Biochemistry 44:13404–13414. https://doi.org/10.1021/bi051008u

10. Mulhbacher J, Lafontaine DA (2007) Ligand recognition determinants of guanine riboswitches. Nucleic Acids Res 35:5568–5580. https://doi.org/10.1093/nar/gkm572

11. Delfosse V, Bouchard P, Bonneau E, et al (2010) Riboswitch structure: an internal residue mimicking the purine ligand. Nucleic Acids Res 38:2057–2068. https://doi.org/10.1093/nar/gkp1080

12. Mandal M, Breaker RR (2004) Adenine riboswitches and gene activation by disruption of a transcription terminator. Nat Struct Mol Biol 11:29–35. https://doi.org/10.1038/nsmb710

13. Pedrolli DB, Kuhm C, Sevin DC, et al (2015) A dual control mechanism synchronizes riboflavin and sulphur metabolism in Bacillus subtilis. Proc Natl Acad Sci U S A 112:14054– 14059. https://doi.org/10.1073/pnas.1515024112

14. Radeck J, Kraft K, Bartels J, et al (2013) The Bacillus BioBrick Box: generation and evaluation of essential genetic building blocks for standardized work with Bacillus subtilis. J Biol Eng 7:29. https://doi.org/10.1186/1754-1611-7-29

15. Zadeh JN, Steenberg CD, Bois JS, et al (2011) NUPACK: Analysis and design of nucleic acid systems. J Comput Chem 32:170–173. https://doi.org/10.1002/jcc.21596

16. Lemay J-F, Desnoyers G, Blouin S, et al (2011) Comparative Study between Transcriptionally - and Translationally-Acting Adenine Riboswitches Reveals Key Differences in Riboswitch Regulatory Mechanisms. PLoS Genet 7:e1001278. https://doi.org/10.1371/journal.pgen.1001278

17. Gong S, Wang Y, Zhang W (2015) Kinetic regulation mechanism of pbuE riboswitch. J Chem Phys 142:15103. https://doi.org/10.1063/1.4905214

18. Pedrolli DB, Matern A, Wang J, et al (2012) A highly specialized flavin mononucleotide riboswitch responds differently to similar ligands and confers roseoflavin resistance to Streptomyces davawensis. Nucleic Acids Res 40:8662–8673. https://doi.org/10.1093/nar/gks616

19. Mandal M, Boese B, Barrick JE, et al (2003) Riboswitches control fundamental biochemical pathways in Bacillus subtilis and other bacteria. Cell 113:577–586

20. Johnson GE, Lalanne J-B, Peters ML, Li G-W (2020) Functionally uncoupled transcription– translation in Bacillus subtilis. Nature 58 5:124–128. https://doi.org/10.1038/s41586-020-2638-5

21. Wickiser JK, Winkler WC, Breaker RR, Crothers DM (2005) The speed of RNA transcription and metabolite binding kinetics operate an FMN riboswitch. Mol Cell 18:49–60. https://doi.org/10.1016/j.molcel. 2005.02.032

22. Bennett BD, Kimball EH, Gao M, et al (2009) Absolute metabolite concentrations and implied enzyme active site occupancy in Escherichia coli. Nat Chem Biol 5:593–599. https://doi.org/10.1038/nchembio.186

23. Kirchner M, Schneider S (2017) Gene expression control by Bacillus anthracis purine riboswitches. RNA 23:762–769. https://doi.org/10.1261/rna.058792.116

