## SUPPLEMENTARY MATERIAL for "Characterization of five purine riboswitches in cellular and cell-free expression systems"

### CONTENT

**Figure S1.** Purine riboswitches apparent affinity ($T_{50}$) determined by *in vitro* gene expression.

**Figure S2**. Growth of *B. subtilis* strains. $OD_{600}$ measurements were taken during growth of *B. subtilis* strains.

**Figure S3**. Relative gene expression under control of the purine riboswitches in *B. subtilis*.

**Figure S4.** Relative gene expression under control of the purine riboswitches in *B. subtilis*.

**Figure S5.** Correlation between the transcription readthrough and the measured reporter expression under control of the purine riboswitches.

**Table S1**. Plasmids used in this study

**Table S2**. DNA parts and oligonucleotides used in this study

**Table S3**. Strains used and generated in this research

**Sequences**

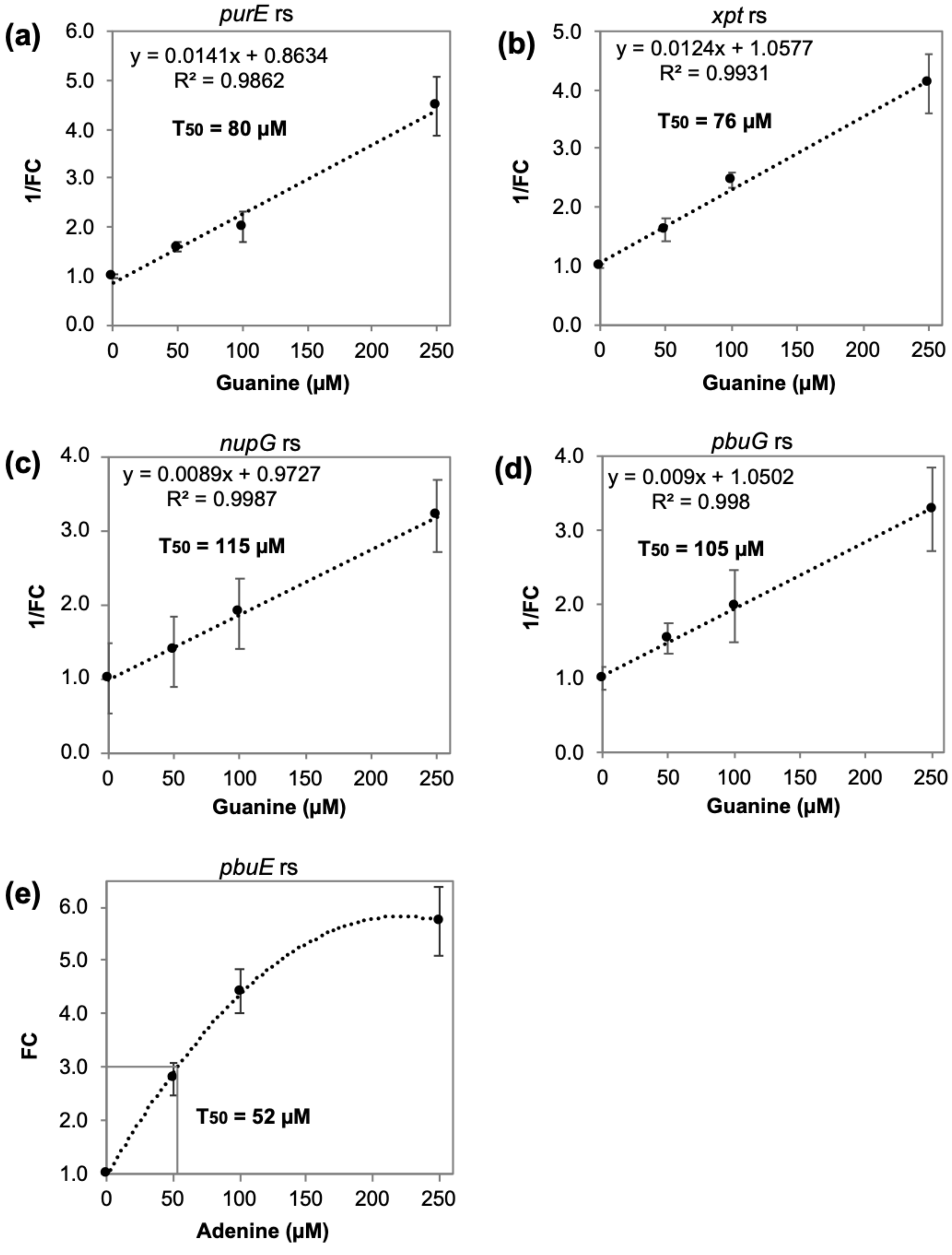


**Fig. S1** Purine riboswitches apparent affinity ($T_{50}$) determined by *in vitro* gene expression. Guanine concentration versus the inverse of the fold change (1/FC) was plotted for the OFF riboswitches (a) *purE*, (b) *xpt*, (c) *nupG*, and (d) pbuG, and (e) adenine concentration versus the fold change (FC) was plotted for the ON riboswitch *pbuE*. All data refers to the in vitro gene expression assay shown in figure 3. Error bars represent the standard deviation calculated from biological triplicates.

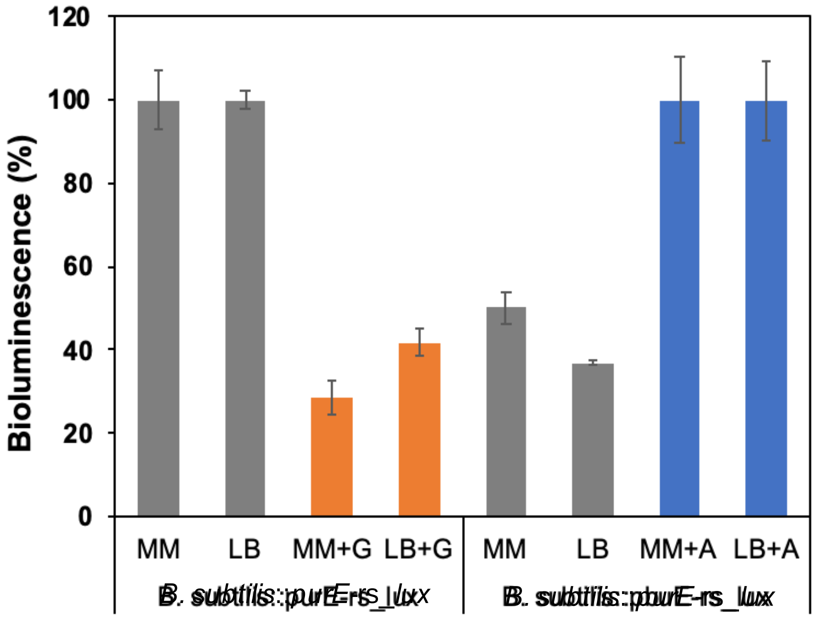


**Fig. S2** Purine riboswitches performance in *B. subtilis* cultivated in mineral medium (MM) or LB medium supplemented with guanine or adenine. The *lux* operon, under control of either the *purE* or the *pbuE* riboswitch, was introduced as a single-copy into *B. subtilis* genome. The resulting strains were cultured for four hours after receiving treatment (100 µM guanine or adenine). Bioluminescence output was normalized to the controls without purine addition.

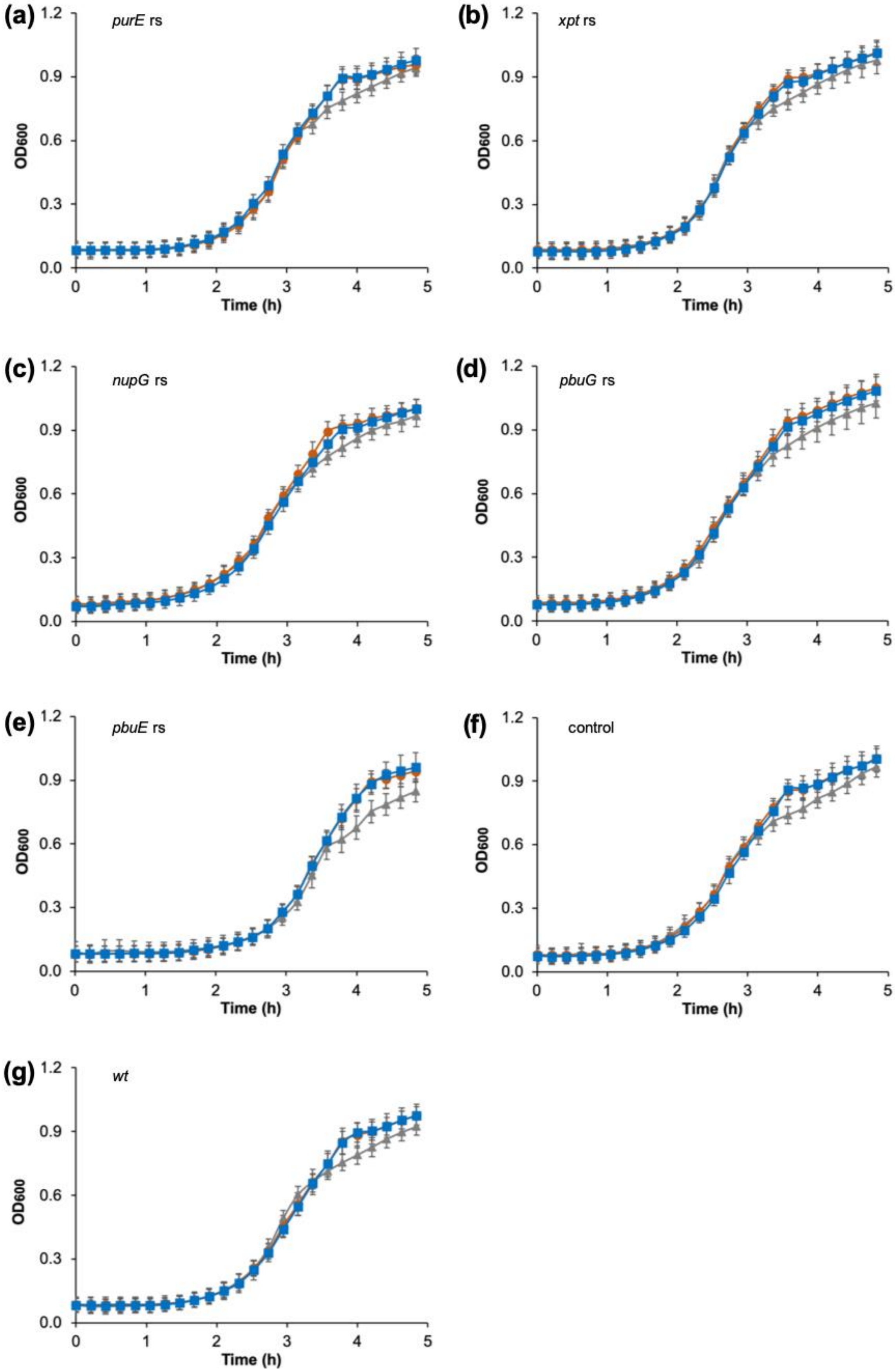
(a)
purE rs
(b)
xpt rs
(c)
nupG rs
(d)
pbuG rs
(e)
pbuE rs
(f)
control
(g)
wt
OD600
Time (h)
0.0
0.3
0.6
0.9
1.2
0
1
2
3
4
5

**Fig. S3** Growth of *B. subtilis* strains. $OD_{600}$ measurements were taken during the growth of *B. subtilis* strains. Three treatments were performed: control - LB only (gray triangles), LB + 100 µM guanine (orange circles), and LB + 100 µM adenine (blue squares). All strains tested carry one genomic copy of *luxABCDE* under control of the (a) *purE* riboswitch; (b) *xpt* riboswitch; (c) *nupG* riboswitch; (d) *pbuG* riboswitch; or the (e) *pbuE* riboswitch. (f) control strain carrying one genomic copy of *luxABCDE* not regulated by a riboswitch. (g) *B. subtilis wt* strain. Error bars represent the standard deviation calculated from biological triplicates.

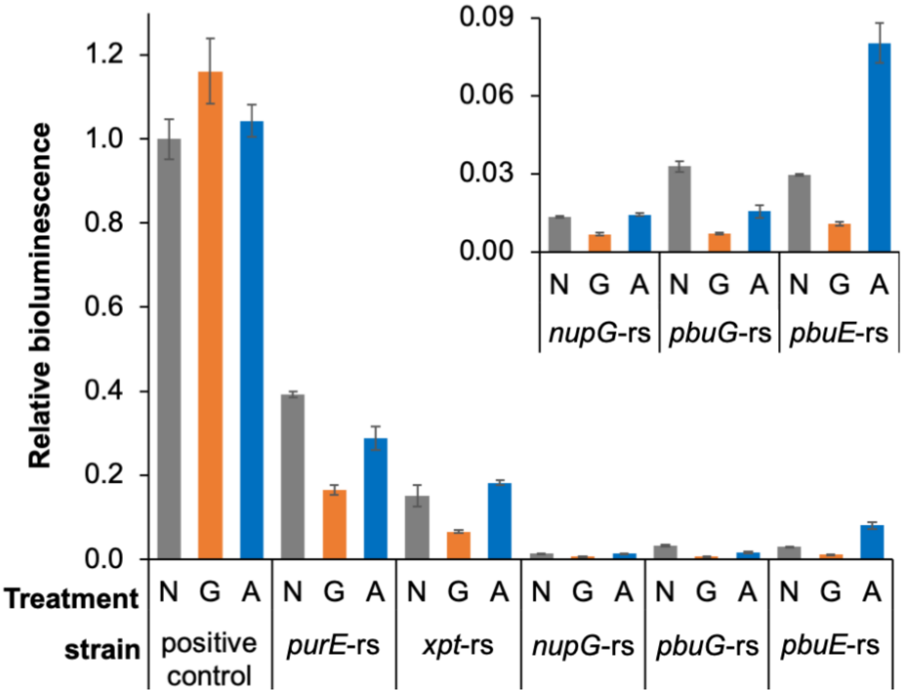


**Fig. S4** Relative gene expression under the control of the purine riboswitches in *B. subtilis*. Bioluminescence levels refer to the maximum bioluminescence measured in the *in vivo* assay shown in Figure 4 for each riboswitch for each treatment: none (N), guanine (G), adenine (A). Bioluminescence output was normalized to the positive control, none treatment level. Error bars represent the standard deviation calculated from biological triplicates.

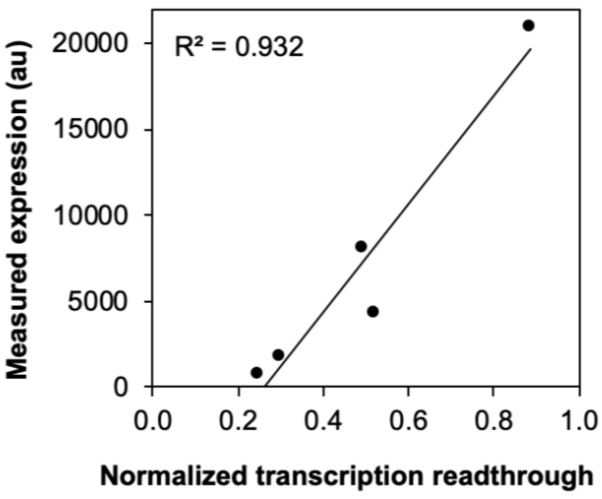


**Fig. S5** Correlation between the transcription readthrough and the measured reporter expression under control of the purine riboswitches. Normalized transcription readthrough refers to the levels full length RNA measured after *in vitro* transcription. Measured expression refers to the maximum bioluminescence measured in the *in vivo* assay shown in figure 4 for control treatments (*pbuG*, *xpt*, *purE*, and *nupG*) or adenine treatment (*pbuE*).

**Table S1**. Plasmids used in this study

| Plasmids | Description | Reference |
|---|---|---|
| **pBS3C*lux*** | Integrates at the *B. subtilis sacA* locus, chloramphenicol resistance (BBa_K823025) | (Radeck et al., 2013) |
| **$P_{ribB}$-*luc*** | The firefly luciferase gene under control of the *E. coli ribB* promoter (sigma 70) | (Pedrolli et al., 2015) |
| **$P_{ribB}$-r*ibG-luc*** | The firefly luciferase gene under control of the *E. coli ribB* promoter and the *ribG* FMN riboswitch of *B. subtilis* | (Pedrolli et al., 2015) |
| **$P_{ribB}$-*purE-luc*** | The firefly luciferase gene under control of the *E. coli ribB* promoter and the *purE* riboswitch of *B. subtilis* | This study |
| **$P_{ribB}$-*xpt-luc*** | The firefly luciferase gene under control of the *E. coli ribB* promoter and the *xpt* riboswitch of *B. subtilis* | This study |
| **$P_{ribB}$-*pbuG-luc*** | The firefly luciferase gene under control of the *E. coli ribB* promoter and the *pbuG* riboswitch of *B. subtilis* | This study |
| **$P_{ribB}$-*nupG-luc*** | The firefly luciferase gene under control of the *E. coli ribB* promoter and the *nupG* riboswitch of *B. subtilis* | This study |
| **$P_{ribB}$-*pbuE-luc*** | The firefly luciferase gene under control of the *E. coli ribB* promoter and the *pbuE* riboswitch of *B. subtilis* | This study |
| **pBS3C-$P_{srfA}$-*lux*** | *luxABCDE* gene cluster under control of the *srfA* promoter of *B. subtilis* | This study |
| **pBS3C-$P_{srfA}$-*purE-lux*** | *luxABCDE* gene cluster under control of the *srfA* promoter and the *purE* riboswitch of *B. subtilis* | This study |
| **pBS3C-$P_{srfA}$-*pbuG-lux*** | *luxABCDE* gene cluster under control of the *srfA* promoter and the *pbuG* riboswitch of *B. subtilis* | This study |
| **pBS3C-$P_{srfA}$-*pbuE-lux*** | *luxABCDE* gene cluster under control of the *srfA* promoter and the *pbuE* riboswitch of *B. subtilis* | This study |
| **pBS3C-$P_{srfA}$-*nupG-lux*** | *luxABCDE* gene cluster under control of the *srfA* promoter and the *nupG* riboswitch of *B. subtilis* | This study |
| **pBS3C-$P_{srfA}$-*xpt-lux*** | *luxABCDE* gene cluster under control of the *srfA* promoter and the *xpt* riboswitch of *B. subtilis* | This study |

**Table S2**. DNA parts and oligonucleotides used in this study

| Parts and oligos | Sequence | Description |
|---|---|---|
| **M13.seq-fw** | TGTAAAACGACGGCCAGT | Sequencing primers |
| **M13.seq-rv** | CAGGAAACAGCTATGAC | |
| **pBS3C*lux*.seq_rv** | GAAATGATGCTCCAGTAACC | |
| **BioBrick-suffix_fw** | TACTAGTAGCGGCCGCTGCAG | Insertion of Biobrick prefix (BBp) and suffix (BBs) |
| **BioBrick-prefix_rv** | CTCTAGAAGCGGCCGCGAATTC | |
| ***purE*(*Hind*III)_fw** | GCTAAAGCTTGAAATCAAAACACGACC TCATATAATC | Oligos used for amplifying the riboswitches for cloning into the pP$_{ribB}$-luc |
| ***purE*(*Bam*HI)_rv** | TCGAGGATCCATTCCTACTAGCGGCTG C | |
| ***xpt*(*Hind*III)_ fw** | GCTAAAGCTTAGGAACACTCATATAAT CGCGTG | |
| ***xpt*(*Bam*HI)_rv** | TCGTGGATCCTGCTTCCATCCTGTCTA C | |
| ***pbuE*(*Hind*III)_fw** | GCTAAAGCTTATTATCACTTGTATAAC CTCAATAATATG | |
| ***pbuE*(*Bam*HI)_rv** | TCGAGGATCCAACAAACTCCTTTACTT AAATG | |
| ***nupG*(*Hind*III)_fw** | GCTAAAGCTTCATCTTAGAAAAAGACA TTCTTG | |
| ***nupG*(*Bam*HI)_rv** | TCGTGGATCCGATAGCAACTCCTATGA G | |
| **P$_{srfA}$(BBp)_fw** | GCTAGAATTCGCGGCCGCTTCTAGGAT TGAACGCAGCAGTTTGG | Oligos used for amplifying parts for cloning into the pBS3Clux |
| **P$_{srfA}$(*Spe*I)_rv** | TCGAACTAGTTTATCCATATCAGCTTT TAATTCTTATG | |
| ***pbuE*(BBp)_fw** | ATCGGAATTCGCGGCCGCTTCTAGAAT TATCACTTGTATAACCTCAATAATATG | |
| ***pbuE*(BBs)_rv** | TCGACTGCAGCGGCCGCTACTAGTAAA CAAACTCCTTTACTTAAATG | |
| ***xpt*(BBp)_fw** | GCTAGAATTCGCGGCCGCTTCTAGAAG GAACACTCATATAATCG | |
| ***xpt*(BBs)_rv** | TCGTCTGCAGCGGCCGCTACTAGTATG CTTCCATCCTGTCTAC | |
| ***nupG*(BBp)_fw** | ATGCGAATTCGCGGCCGCTTCTAGACA TCTTAGAAAAAGACATTC | |
| ***nupG*(BBs)_rv** | TCGTCTGCAGCGGCCGCTACTAGTAGA TAGCAACTCCTATGAG | |
| ***purE*(BBp)_fw** | TGCAGAATTCGCGGCCGCTTCTAGGGA GTTCTGAGAATTGGTATGC | |
| ***purE*(BBs)_rv** | ATCGCTGCAGCGGCCGCTACTAGTATC GAGGATCTCTGTTCCCCAC | |
| ***PT7-RS_fw*** | TAATACGACTCACTATAGGGGATGAAG CGTTATAGTGAATCCGCTTAAAGC | Oligos used to PCR amplify riboswitch linear templates for in *vitro transcription* |
| ***luc164_rv*** | CGATCGTACGTGATGTTCACCTCGATA TGTGC | |

**Table S3**. Strains used and generated in this research

| Strain | Description | Reference |
|---|---|---|
| ***E. coli* Top 10** | Cloning strain | Invitrogen |
| ***B. subtilis* 168** | Wild-type, *trpC2* | Lab collection |
| ***B. subtilis*::P$_{srfA}$-*luxABCDE*** | P$_{srfA}$-*luxABCDE* integrated into the *sacA locus* | This study |
| ***B. subtilis*::*luxABCDE*** | *luxABCDE* integrated into *sacA locus* | This study |
| ***B. subtilis*::P$_{srfA}$-*purE*-*luxABCDE*** | P$_{srfA}$-*purE*-*luxABCDE* integrated into the *sacA locus* | This study |
| ***B. subtilis*::P$_{srfA}$-*pbuE*-*luxABCDE*** | P$_{srfA}$-*pbuE*-*luxABCDE* integrated into the *sacA locus* | This study |
| ***B. subtilis*::P$_{srfA}$-*nupG*-*luxABCDE*** | P$_{srfA}$-*nupG*-*luxABCDE* integrated into the *sacA locus* | This study |
| ***B. subtilis*::P$_{srfA}$-*pbuG*-*luxABCDE*** | P$_{srfA}$-*pbuG*-*luxABCDE* integrated into the *sacA locus* | This study |
| ***B. subtilis*::P$_{srfA}$-*xpt*-*luxABCDE*** | P$_{srfA}$-*xpt*-*luxABCDE* integrated into the *sacA locus* | This study |

### Sequences

#### 1. Riboswitches

***purE***
GCTAAAGCTTGAAATCAAAACACGACCTCATATAATCTTGGGAATATGGCCCATAAGTTTCTACCCGGCAACCGT
AAATTGCCGGACTATGCAGGAAAGTGATCGATAAAACTGACATGGATATATCGCAGAAGCGAACGACTGACGATA
CATGTACCATGCCCGGTTTGTATTGCTTCCTCATAAGTGCAATGCAGAGCGGGTATTTTTTATTTTCTGAAAACA
AAAGCATTAGAAGGTGGGGAACAGAGGATCCTCGA

***nupG***
CATCTTAGAAAAAGACATTCTTGTATATGATCAGTAATATGGTCTGATTGTTTCTACCTAGTAACCGT
AAAAAACTAGACTACAAGAAAGTTTGAATAAATTTGAACGAGTTGAAAAGGACAAGTTCTTTTCTGTT
TGCTCTTATTTTTCACACTTTCTGCACTTCCAGACTTTGTGAAGGATAAGAGCTTTTTTTGTTTCCAT
AATAACCCTCATAGGAGTTGCTATC

***pbuE***
ATTATCACTTGTATAACCTCAATAATATGGTTTGAGGGTGTCTACCAGGAACCGTAAAATCCTGATTA
CAAAATTTGTTTATGACATTTTTTGTAATCAGGATTTTTTTTATTTATCAAAACATTTAAGTAAAGGA
GTTTGTT

***pbuG***
ACACAGAAATCAAATAAGATGAATTCGTATAATCGCGGGAATATGGCTCGCAAGTCTCTACCAAGCTA
CCGTAAATGGCTTGACTACGTAAACATTTCTTTCGTTTGATATAAATAAAACACGGTTATTTATTCAA
ACTGAAATCCGTCTGTAGTCAAGCGTCCCAAAATGTATTGGGACGTTTTTTATTTGGCGTTTTCAGGA
CAGAATAAATAGCAAAGAGTGAAGGGAGTCAAATAGC

***xpt***
AGGAACACTCATATAATCGCGTGGATATGGCACGCAAGTTTCTACCGGGCACCGTAAATGTCCGACTA
TGGGTGAGCAATGGAACCGCACGTGTACGGTTTTTTGTGATATCAGCATTGCTTGCTCTTTATTTGAG
CGGGCAATGCTTTTTTTATTCTCATAACGGAGGTAGACAGGATGGAAGCA

***ribG***
TTGTATCCTTCGGGGCAGGGTGGAAATCCCGACCGGCGGTAGTAAAGCACATTTGCTTTAGAGCCCGT
GACCCGTGTGCATAAGCACGCGGTGGATTCAGTTTAAGCTGAAGCCGACAGTGAAAGTCTGGATGGGA
GAAGGATGATGAGCCGCTATGCAAAATGTTTAAAAATGCATAGTGTTATTTCCTATTGCGTAAAATAC
CTAAAGCCCCGAATTTTTTATAAATTCGGGGCTTTTTTGACGGTAAATAACAAAA

#### 2. Promoters

$\mathbf{P}_{ribB}$
GGCG**TTGCTG**CCGCTAATCATTAGCGT**TATAGT**GAATCCGCTTA

$\mathbf{P}_{srfA}$
GATTGAACGCAGCAGTTTGGTTTAAAAATTTTTATTTTTCTGTAAATAATGTTTAGTGGAAATGATTG
CGGCATCCCGCAAAAAATATTGCTGTAAATAAACTGGAATCTTTCGGCATCCCGCATGAAACTTTTCA
CCCATTTTTCGGTGATAAAAACATTTTTTTCATTTAAACTGAACGGTAGAAAGATAAAAAATATTGAA
AACAATGAATAAATAGCCAAAATTGGTTTCTTATTAGGGTGGGGTCTTGCGGTCTTTATCCGCTTATG
TTAAACGCCGCAATGCTGACTGACGGCAGCCTGCTTTAATAGCGGCCATCTGTTTTTTGA**TTGGAA**GC
ACTGCTTTTTAAGTG**TAGTACT**TTGGGCTATTTCGGCTGTTAGTTCATAAGAATTAAAAGCTGATATG
GATAA

**Features:**
-35 promoter box
-10 promoter box

### REFERENCES

Pedrolli, D.B., Kuhm, C., Sevin, D.C., Vockenhuber, M.P., Sauer, U., Suess, B., Mack, M., 2015. A dual control mechanism synchronizes riboflavin and sulphur metabolism in Bacillus subtilis. Proc Natl Acad Sci U S A 112, 14054–14059. https://doi.org/10.1073/pnas.1515024112

Radeck, J., Kraft, K., Bartels, J., Cikovic, T., Durr, F., Emenegger, J., Kelterborn, S., Sauer, C., Fritz, G., Gebhard, S., Mascher, T., 2013. The Bacillus BioBrick Box: generation and evaluation of essential genetic building blocks for standardized work with Bacillus subtilis. J. Biol. Eng. 7, 29. https://doi.org/10.1186/1754-1611-7-29
